# N-terminal cysteine independent effects of oxidative stress on the stability of RGS4 and 5

**DOI:** 10.64898/2026.04.20.719627

**Authors:** Haeun Kim, Nianhan Shang, Ya-Min Tian, Peter J Ratcliffe, Thomas P Keeley

**Author notes:** Correspondence should be addressed to: Thomas P. Keeley, NDM Research Building, University of Oxford, Oxford OX3 7FZ, UK 01865613960. These authors contributed equally and are listed alphabetically (Haeun Kim, Nianhan Shang, Ya-Min Tian).

## Abstract

Oxidative post-translational modifications on the sulfhydryl group of cysteines can occur spontaneously or enzymatically. The enzymatic dioxygenation of N-terminal cysteines has emerged as a new oxygen sensing paradigm, catalysed by 2-aminoethanethiol dioxygenase (ADO) in mammals. Conflicting evidence has been reported in recent years on whether this reaction can occur spontaneously in the absence of ADO. Here we sought to address whether cellular oxidative stress can interfere with ADO-catalysed N-terminal cysteine dioxygenation. Using a system to produce titratable intracellular levels of H_2_O_2_, we demonstrate that the stability of RGS4 and 5 is not affected by physiological levels of oxidative stress, whether ADO is present or not. However, cytotoxic levels of oxidative stress induced by tert-butyl hydroperoxide did induce a substantial increase in RGS4/5 protein stability that occurred independently of the Cys N-degron pathway. This effect of tBHP was abolished by Fe^2+^ chelation, lysosomal pH perturbation, and lipid peroxide scavenging. We conclude that N-terminal cysteine dependent proteolysis of RGS4/5 is not sensitive to physiological oxidative stress, but these proteins can be stabilised during the process of oxidative stress-induced cell death through an N-terminal cysteine independent mechanism.

## Introduction

Cysteine undergoes a variety of post-translational modifications and distinguishing whether these occur enzymatically or non-enzymatically continues to be an important problem. One of the most common forms of cysteine modification is sulfhydryl oxidation to form cysteine sulfenic (-SOH), sulfinic (-SO_2_H) and sulfonic (-SO_3_H) acids. Whilst -SOH modifications are highly prevalent and can act as reversible signalling events^1^, more complete oxidation to -SO_2_H and -SO_3_H is relatively rarer and under physiological conditions likely confined to enzyme-coupled reactions or non-signalling accumulative modifications^2^. Examples of enzyme-coupled cysteine sulfinylation include those involving peroxiredoxins^3^, DJ-1^4^ and N-terminal cysteine dioxygenation^5^. The latter has emerged as an important process in eukaryotic oxygen sensing, wherein proteins bearing an N-terminal cysteine are subjected to rapid dioxygenation (to Cys-SO_2_H) and degradation by the N-degron pathway with extreme sensitivity to O_2_ availability^6^. In mammals this reaction targets the regulators of G-protein signalling (RGS) 4, 5, 16 and interleukin-32 (IL-32), and is catalysed by 2-aminoethanethiol dioxygenase (ADO)^7^. N-terminal cysteine-SO_2_H containing proteins are then arginylated by an arginyl transferase (ATE1), ubiquitinylated by UBR box proteins and degraded by the 26S proteasome^6^. Thus, when O_2_ levels are insufficient to support ADO activity, substrate proteins will rapidly accumulate within the cell. Increased stability of RGS4/5 through genetic inhibition of ADO or ATE1 has been linked to cellular sensitivity to Gαq-protein coupled receptor activation^7^, though the physiological processes affected by this action remain unexplored.

In recent years contrasting evidence has been presented concerning the propensity of N-terminal cysteine dioxygenation to occur non-enzymatically via the spontaneous action of reactive oxygen or nitrogen species^8-10^. As interactions between oxygen sensing pathways and oxidative stress have long been appreciated^11-14^, we set out to establish whether or not oxidative stress can influence the stability of ADO substrate proteins. We considered two pathways through which reactive oxygen species might influence the stability of ADO substrates non-enzymatically; either single oxidation of the N-terminal cysteine to sulfenic acid, or double oxidation to N-terminal cysteine sulfinic acid (Fig. 1A). Importantly, single oxidation does not create a substrate for ATE1-mediated N-degron proteolysis thus would be predicted to lead to stabilisation, whereas the double oxidation would bypass the reaction catalysed by ADO leading to persistent instability even under hypoxic conditions.

**Figure 1.**
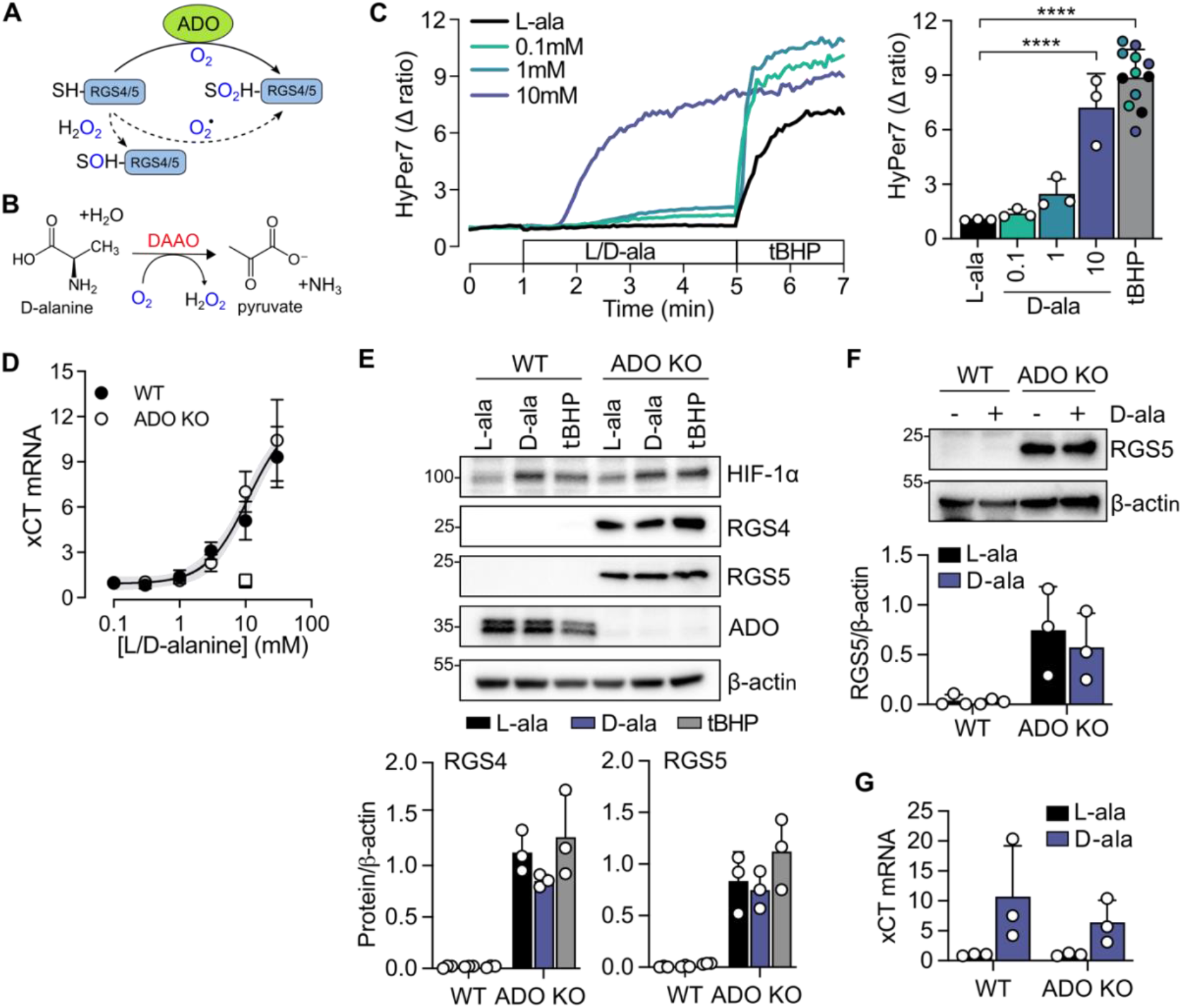
Physiological levels of oxidative stress do not impact RGS4/5 protein stability. (**A**) A schematic illustrating ADO-catalysed dioxygenation of N-terminal cysteine residues in RGS4 and RGS5 (solid line) and potential alternative, non-enzymatic oxidation pathways (dashed lines). (**B**) Mechanism of action of D-amino acid oxidase (DAAO), used to produce intracellular H_2_O_2_. (**C**) SH-SY5Y cells expressing DAAO were exposed to different concentrations of D-alanine and H_2_O_2_ production monitored using the H_2_O_2_ sensor HyPer7. 10mM L-alanine was used as a negative control, and 10μM tBHP was applied at the end of each experiment for comparison and as a positive control. A representative trace is provided on the left, and the average change in HyPer7 ratio at 5 minutes (for L/D-ala) and 7 minutes (for tBHP) for each experiment is plotted on the accompanying histogram. ****P<0.001 vs L-ala. (**D**) WT or ADO KO SH-SY5Y cells expressing DAAO and treated with increasing concentrations of D-alanine, or 10mM L-alanine as a control (squares) for 16 hours, and xCT mRNA measured by RT-qPCR and expressed relative to HPRT mRNA. Least squares regression analysis confirmed one curve can accurately represent both datasets (p=0.09, F=2.169 [4, 41], extra sum-of-squares F test). (**E**) Cells were treated with L/D-alanine (10mM) or 10μM tBHP for 4 hours, and then levels of RGS4 and 5 protein assessed by immunoblotting. HIF-1α protein levels were as a comparison. (**F**) Similar to E, but cells were treated with 10mM L/D-alanine for 16h. Cell extracts were immunoblotted to determine RGS5 protein levels, and (**G**) xCT mRNA levels were assessed by RT-qPCR to serve as a positive control for oxidative stress. All immunoblots are representative of at least 3 replicate experiments, and all data are presented as the mean ± S.D. from at least 3 independent experiments.

## Results

To induce physiological levels of oxidative stress in cells, we employed a D-amino acid oxidase (DAAO) system that enables controlled and titratable intracellular H_2_O_2_ production (Fig. 1B)^15-17^. Through co-expression with the fluorescent H_2_O_2_ reporter HyPer7^18^, this system was confirmed to produce intracellular H_2_O_2_ proportionally and specifically in response to D-alanine (Fig. 1C). The level of intracellular H_2_O_2_ observed in response to 10mM D-alanine was comparable to treatment with 10μM tert-butyl hydroperoxide (tBHP, Fig. 1C). For the purposes of this report, this level of oxidative stress will be defined as physiological because it was associated with no visible change in viability. To examine whether this was sufficient to induce an antioxidant response, we monitored the induction of xCT (SLC7A11) mRNA, the light subunit of the cystine/glutamate antiporter and established redox-sensitive Nrf2-dependent transcript^19^. In both ADO-competent and -deficient SH-SY5Y cells stably expressing DAAO, supplementing the medium with increasing concentrations of D-alanine resulted in dose-dependent induction of xCT (Fig. 1D). No difference was detected between ADO-competent and -deficient cells, suggesting that loss of ADO does not cause a major disturbance in cellular redox homeostasis. Treating SH-SY5Y cells with D-alanine (10mM) or tBHP (10μM) for 4 hours had no detectable impact on RGS4 and 5 protein levels at baseline, nor was any effect evident in ADO-deficient SH-SY5Y cells in which RGS4/5 is constitutively stabilised (Fig. 1E). Notably, these treatments did cause some normoxic stabilisation of HIF-1α, consistent with inhibition of prolyl hydroxylation through direct oxygenase post-translational modification^12^. Prolonged treatment with D-alanine (16 hours) still had no detectable impact on RGS5 protein levels (Fig. 1F), despite significant induction of xCT mRNA (Fig. 1G).

Exposure of SH-SY5Y cells to higher levels of tBHP (25μM) caused significant loss of cell viability and cell-rounding consistent with programmed cell death, and is defined hereafter as cytotoxic oxidative stress. Surprisingly, we observed remarkably high levels of RGS4 and 5 protein following 4-hour treatment with 25μM tBHP in the remaining viable cells, irrespective of cellular ADO status (Fig. 2A). This was not due to an increase in transcription, as no significant difference in RGS4/5 mRNA expression was observed in cells treated with tBHP (Fig. 2B). To test the generality of these observations, we repeated these experiments in the colon adenocarcinoma cell line, RKO, which expresses the ADO substrates RGS4 and IL-32^7,9^. Notably, cytotoxicity was only observed at much higher concentrations of tBHP (100μM) in these cells. Nonetheless, treatment of ADO-competent or -deficient RKO cells with this higher level of tBHP induced a similar effect on RGS4 protein levels as observed in SH-SY5Y cells, demonstrating concordance between cytotoxic tBHP exposure and RGS4/5 induction. However, IL-32 protein was unaffected by cytotoxic concentrations of tBHP despite containing a cysteine N-degron (Fig. 2C).

**Figure 2.**
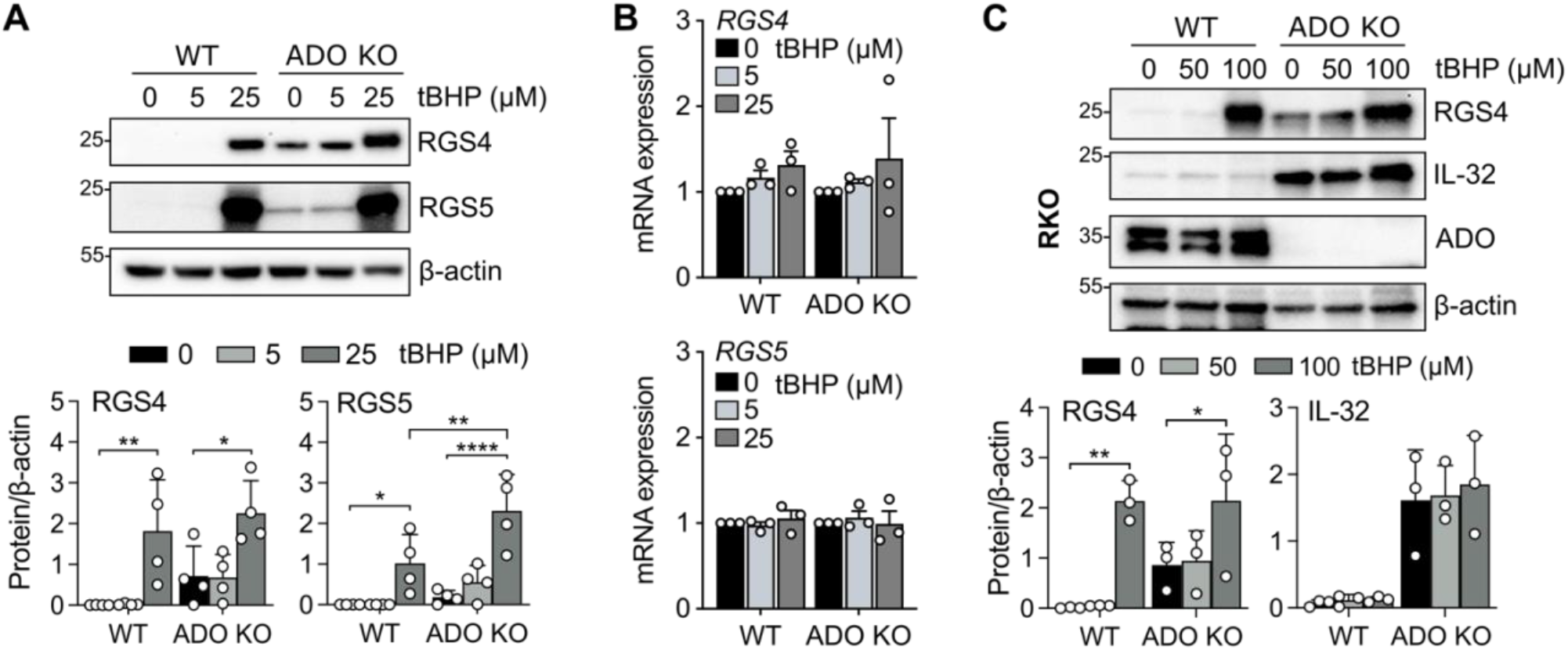
Cytotoxic oxidative stress aberrantly affects RGS4 and 5 but not IL-32. (**A**) Wild-type or ADO deficient SH-SY5Y cells treated with either 5 or 25μM tBHP for 4 hours, and samples immunoblotted for RGS4 and RGS5. (**B**) Parallel extracts were analysed for RGS4 and 5 mRNA levels by RT-qPCR, normalised to HPRT mRNA. (**C**) RKO wild-type or ADO deficient cells were treated with either 50 or 100μM tBHP (causing equivalent cytotoxicity as observed in SH-SY5Y treated with 25μM tBHP) and analysed for RGS4 and IL-32 protein abundance. All immunoblots are representative of at least 3 independent replicates, and all data are presented as the mean ± S.D. from at least 3 independent experiments. *P<0.05, **P<0.01, ****P<0.01, two-way ANOVA with Holm-Sidak post-hoc test.

In the absence of ATE1, ADO substrates exist in either N-terminal cysteine sulfinic (-SO_2_H) or sulfonic acid (-SO_3_H) forms^20^, both of which are constitutively stable. Thus further oxidation, if it can occur, would not be expected to affect stability via the N-degron pathway (Fig. 3A). Despite this, 25μM tBHP still caused a robust increase in RGS5 protein in SH-SY5Y cells deficient in ATE1 (Fig. 3B). In contrast, 2,2-dipyridyl (2,2DIP, an Fe^2+^ chelator that potently inhibits ADO activity^7^) had no effect on the stability of RGS5 in ATE1 KO cells, confirming disruption of the Cys N-degron pathway in these cells.

**Figure 3.**
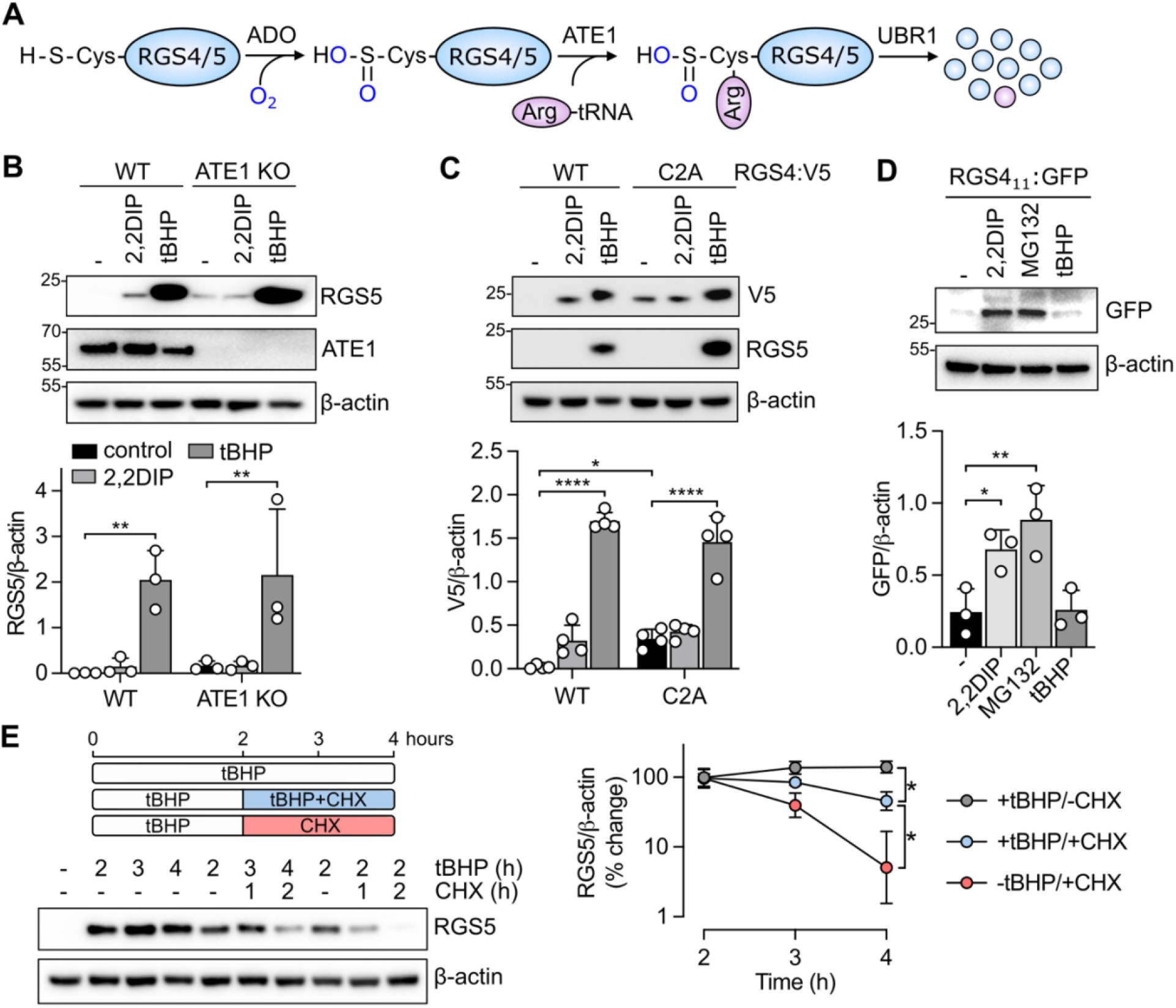
tBHP acts on RGS4/5 stability through an N-terminal cysteine independent mechanism. **(A)** The actions of ADO and ATE1 on the Cys N-degron pathway. (**B**) WT or ATE1-deficient SH-SY5Y cells were treated with either 100μM 2,2 dipyridyl (2,2DIP, an iron chelator) or 25μM tBHP for 4 hours. RGS5 protein adundance was assayed by immunoblotting, with ATE1 immunoblotted as validation of the knock-out. (**C**) Cells stably expressing either WT RGS4 or a C2A mutant variant, both C-terminally tagged with V5 to discriminate from endogenous RGS4, were treated with 100μM 2,2DIP or 25μM tBHP for 4 hours, then adundance of the exogenous RGS4 protein assessed by immunoblotting using a V5 antibody. Endogenous RGS5 was provided as an internal control. (**D**) Cells stably expressing a fusion reporter protein, comprised of the first 11 amino acids of RGS4 appended to eGFP, were treated with 2,2DIP (100μM), MG132 (25μM) or tBHP (25μM) for 4 hours. Accumulation of the reporter protein was assessed by immunoblotting for GFP. (**E**) Cells were treated with tBHP and/or cycloheximide (CHX, 10μM) as indicated in the schematic, then extracts blotted for RGS5 adundance. All immunoblots are representative of at least 3 independent replicates, and all data are presented as the mean ± S.D. from at least 3 independent experiments. *P<0.05, **P<0.01, ****P<0.0001, one/two-way ANOVA with Holm-Sidak post-hoc test.

Based on this data, we hypothesised that tBHP may exert its effect on RGS4/5 through a mechanism independent of the N-terminal cysteine. To test this, we expressed C-terminal V5-tagged RGS4 with either its native cysteine (WT) or a mutant in which this cysteine is replaced with alanine (C2A), in SH-SY5Y cells. As shown in Figure 3C, only the WT RGS4:V5 protein showed ADO-dependent regulation, as evidenced by an increase in protein level when treated with 2,2DIP. In contrast, both the WT and C2A RGS4:V5 variants demonstrated considerable increases in detectable protein following treatment with cytotoxic levels of tBHP. To further investigate the apparent lack of dependence of this phenomenon on the N-terminal sequence, we used an established Cys N-degron reporter system in which the first 11 amino acids of RGS4 are appended to the N-terminus of GFP, conferring ADO-dependent instability on the fluorescent protein^21^. This construct allows for the isolation of ADO/N-degron pathway activity on the N-terminus from factors impinging on other parts of the molecule. When expressed in SH-SY5Y cells, RGS4_1-11_GFP was stabilised by 2,2DIP and MG132 (a proteasomal inhibitor), but was unaffected by cytotoxic levels of tBHP (Fig. 3D), further suggesting tBHP-induced RGS4/5 accumulation is independent of the N-terminal cysteine.

We next sought to determine whether the observed effects of tBHP on RGS4/5 levels resulted from a change in protein stability. Cells were treated with tBHP for 2 hours to allow for the accumulation of RGS5 protein, followed by a further 1 or 2 hours incubation with tBHP, or with cycloheximide (a protein synthesis inhibitor) in the presence or absence of tBHP (Fig. 3E). Treatment with cycloheximide resulted in a gradual loss of RGS5 protein over the next 2 hours that was significantly accelerated in the absence of tBHP (estimated t_1/2_ 3.9 vs 2.9 hours). Notably, both half-lifes are considerably longer than the maximum half-life of RGS5 we have previously determined (approximately 4 minutes^9^).

We hypothesised that the observed effects of high concentrations of tBHP on RGS4/5 might be a result of cells undergoing cell death. To test this hypothesis, we treated SH-SY5Y cells with either menadione (an oxidative stress-inducing compound), ICL670A (an Fe^2+^ chelator that does not affect ADO activity^9^) or ionomycin (inducing mitochondrial Ca^2+^ overload^22^) at a concentration titrated to achieve cytotoxicity equivalent to that induced by 25μM tBHP (Fig. 4A). However, none of these compounds had any substantial impact on RGS5 protein levels comparable to tBHP (Fig. 4B). It has been proposed that cytotoxicity arising from exposure to tBHP/H_2_O_2_ arises primarily from the Fe^2+^ oxidation to Fe^3+^, accompanied by the generation of the more oxidising OH^•^ radical. This reaction occurs predominantly within the lysosome, in which a substantial pool of redox-active free iron accumulates due to the breakdown of iron-containing proteins^23^. Accordingly, chelation of intracellular Fe^2+^ using desferoxamine (DFO) has been shown to protect cells against ROS-dependent cell death^24,25^. Importantly, DFO does not inhibit ADO activity unlike 2,2DIP^9^. Consistent with previous findings, we observed a similar cytoprotection in SH-SY5Y cells here (Fig. 4C) which was accompanied by a substantial reduction in RGS5 protein levels (Fig. 4D).

**Figure 4.**
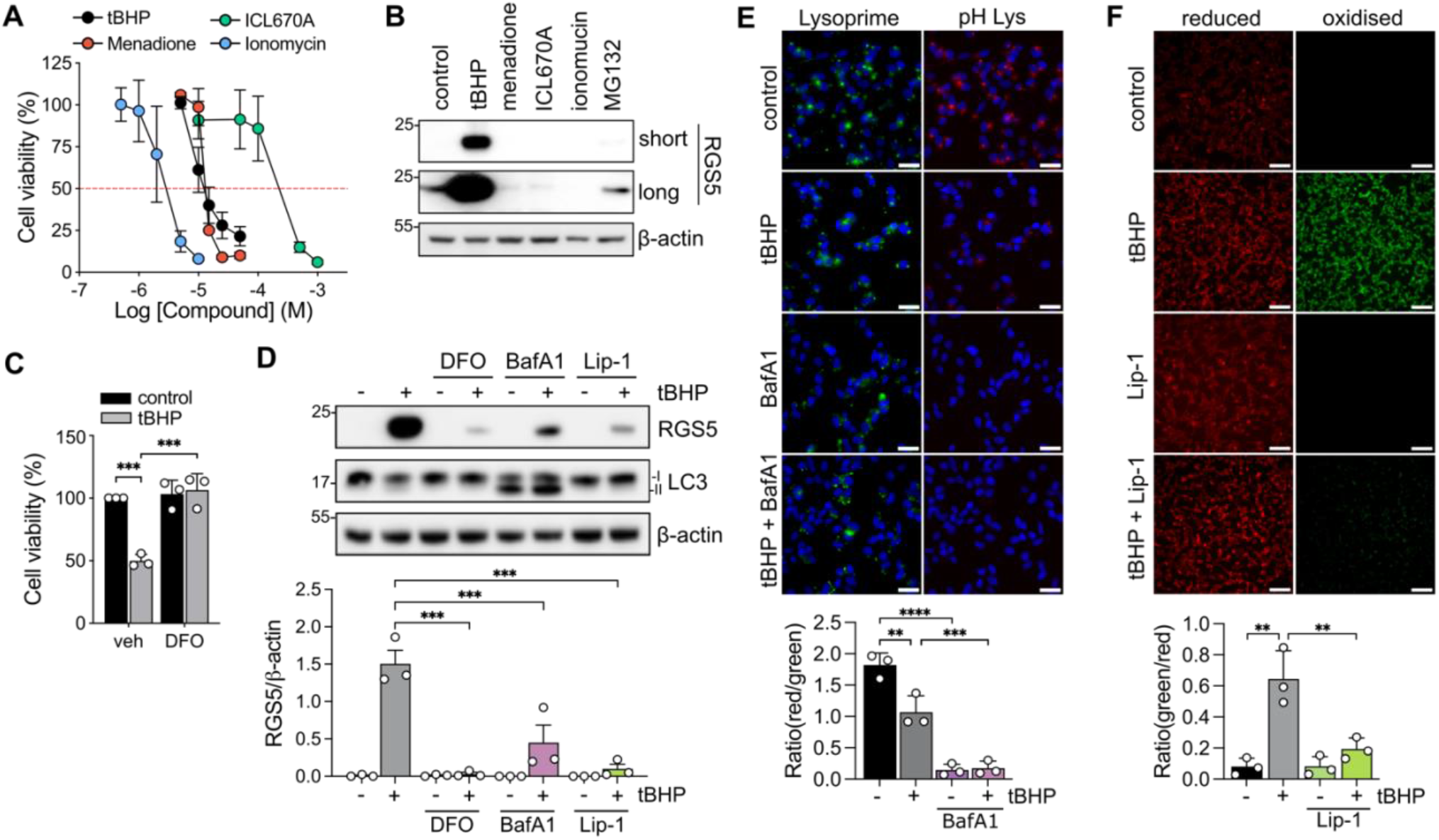
Oxidative damage in the lysosome promotes RGS4/5 stability. (**A**) SH-SY5Y cells were treated with the indicated concentrations of tBHP, menadione (an oxidative stress-inducing compound), ICL670A (a cytotoxic Fe^2+^-chelator) or ionomycin (to cause Ca^2+^-dependent cell death), and viability assessed 4 hours later by MTT staining. (**B**) Cells were treated with a concentration of each compound chosen to elicit equivalent cell death to 25μM tBHP (as indicated by the dashed red line, 1.5μM ionomycin, 200μM ICL670A, 12.5μM menadione), then the accumulation of RGS5 assessed by immunoblotting. Short and long refer to different exposure times of the same membrane. (**C**) Cells were treated with 25μM tBHP in the presence or absence of 100μM DFO (an iron chelator) for 4 hours, and viability assessed by MTT staining. (**D**) Cells were treated for 4 hours with 25μM tBHP, DFO (100μM), bafilomycin A1 (BafA1, 200nM 16h pretreatment) or liproxstatin-1 (Lip-1, 1μM) alone or in combination as indicated. RGS5 protein abundance was assessed by immunoblotting, and LC3-II protein abundance were also assessed as a positive control for lysosomal inhibition. (**E**) Cells were treated for 4 hours with 25μM tBHP, 200nM BafA1 or both and then stained with Lysoprime green (a pH-insensitive lysosome stain) and pH Lys red (a pH-sensitive lysome stain), and fluorescence assessed. Representative images at both excitation wavelenghts are provided, and the quantification presented as a ratio of red/green fluorescence from 3 independent experiments presented below. Scale bar is 30μm. (**F**) Cells were treated for 4 hours with 25μM tBHP, 1μM Lip-1 (a ferroptsis inhibitor) or both and lipid peroxidation assessed using BODIPY-C11, which exhibits red and green fluorescence when reduced/oxidised, respectively. Representative images at both excitation wavelengths are provided, and the quantification presented as a ratio of green/red fluorescence from 3 independent experiments presented below. Scale bar is 100μm. All immunoblots are representative of at least 3 independent replicates, and all data are presented as the mean ± S.D. from at least 3 independent experiments. **P<0.01, ***P<0.001, ****P<0.0001, one/two-way ANOVA with Holm-Sidak post-hoc test.

Fenton reactions between H_2_O_2_ and Fe^2+^ also require an acidic pH, and thus we reasoned that perturbations in lysosomal pH imposed by inhibiting the vacuolar ATPase using bafilomycin A1 (BafA1) could attenuate the effect of tBHP on RGS5 stability. Indeed, treatment of SH-SY5Y cells with BafA1 dissipated the lysosomal pH gradient (Fig. 4E) and attenuated tBHP-induced RGS5 accumulation (Fig. 4D). Notably, BafA1 alone did not increase RGS5 despite significant impairment of autophagic degradation, as evidenced by the accumulation of LC3-II (Fig. 4D). Fe^2+^-dependent cell death usually results from an accumulation of lipid perioxidation products^26^. Indeed, radical-trapping antioxidants acting specifically within phospholipid bilayers, such as liproxstatin-1, can effectively prevent Fe^2+^-dependent cell death^27,28^. In line with this, liproxstatin-1 (1μM) potently reduced lipid peroxidation detected using BODIPY-C11 (Fig. 4F) and attenuated RGS5 protein accumulation in response to tBHP (Fig. 4D).

Together, these findings suggest the tBHP-induced RGS4/5 accumulation is likely to arise from iron-dependent redox stress originating within the lysosomal compartment, rather than generalised cell death or lysosomal stress in isolation.

## Discussion

In this work we have addressed the question as to whether physiological cellular oxidative stress can impact ADO-catalysed N-degron proteolysis. This question is of high importance given the susceptibility of cysteine residues to oxidative modifications, as well as the complex interplay between proposed oxygen sensing pathways and reactive oxygen species. Using a well-established system to generate consistent and titratable levels of intracellular H_2_O_2_, we have demonstrated that physiological non-cytotoxic levels of oxidative stress do not impact on the steady-state levels of ADO substrate proteins. In contrast, another proteolytic oxygen sensing pathway operating in mammals, the PHD-HIFα pathway, is susceptible to oxidative modifications at multiple levels^11-14^. This difference is consistent with our previous work comparing the two pathways^9^, strengthening our hypothesis that ADO-mediated oxygen sensing operates almost exclusively as a molecular oxygen sensing pathway^9,10^.

Our findings contrast with those reported by Heo et al.^8^, which suggested that oxidative stress can oxidise the N-terminal cysteine of RGS4/5 independently of ADO, resulting in degradation via the lysosome. Both that of Heo et al.^8^ and our work highlight a critical role of the lysosome in regulating levels of RGS4 and 5 in response to oxidative stress, yet with opposing effects of bafilomycin A1 and conflicting evidence for the involvement of the N-terminal cysteine. Whilst we cannot fully explain these discrepancies, we feel it important to point out that the study by Heo et al. used 250 μM tBHP to induce this interaction. This is approximately ten-fold higher than the LC50 concentration observed here (Fig. 4A) and elsewhere^14^, and even more in excess of the maximum levels of oxidants expected in vivo^29^. We could not directly verify their claim of non-enzymatic oxidation of the N-terminal cysteines of RGS4 and 5 using mass spectrometry because neither protein was stabilised sufficiently by non-cytotoxic levels of oxidative stress to permit examination. Instead, we interpret the lack of stabilisation as *ipso facto* evidence of a lack of physiologically relevant oxidation which, if present, would increase stability and lead to protein accumulation. In our experiments, treatment with cytotoxic levels of tBHP (4h, 25μM) resulted in profound increases in RGS4/5 protein abundance in the remaining viable cells. This level of accumulation is consistent with the approximatley 60-fold increase in RGS5 half-life following tBHP treatment relative to the maximal half-life reported previously^9^ (4 min vs ∼3.9h). This is not due to an action on the N-terminal cysteine because (i) it was not observed on the other (non-RGS) ADO substrate IL-32 (Fig. 2C), (ii) it persisted in ATE1-deficient cells, in which the N-terminal cysteine is already oxidised (Fig. 3B), and (iii) it occurred on a transgenic C2A RGS4 mutant protein (Fig. 3C), but not on an artificial ADO substrate reporter bearing the equivalent N-terminal peptide (RGS4_1-11_GFP, Fig. 3D).

Instead, our data suggest that lysosomal redox stress induced by tBHP and potentiated by labile Fe^2+^ contributes to the large increase in RGS4/5 stability (Fig. 4). Many aspects of this pathway bear similarities to the process of ferroptosis, a programmed cell death mediated by iron-dependent lipid peroxidation and membrane dysfunction^30^. Indeed, a designated ferroptosis inhibitor, liproxstatin-1, was found to be very effective in reducing the accumulation of RGS5 in response to cytotoxic tBHP treatment (Fig. 4). Treatment with cytotoxic levels of tBHP may also induce ferroptosis, yet how this might result in the large accumulation of RGS4/5 in ferroptosing cells remains unclear. Accelerated degradation of RGS5 when tBHP was removed (Fig. 3E) implies a partly reversible action of tBHP on the stability of RGS4/5 proteins. RGS4 and 5 are cysteine-rich proteins, with many proposed to undergo acyl modification to modulate protein function and/or localisation^31,32^. These acylated cysteines may be oxidised in response to cytotoxic tBHP treatment^26^, perhaps affecting subcellular localisation in a manner that impacts on their stability. However, these observations are clearly confounded by the frank cytotoxicity and as such their physiological relevance remains unclear and should be interpreted with caution. Whilst more work is required to fully understand the observations reported here and their importance in the mechanistic cascade of ferroptosis, we conclude that N-terminal cysteine dependent proteolysis is not sensitive to oxidative stress within a physiologically relevant range.

## Materials and Methods

### Cell culture and treatments

SH-SY5Y cells were cultured in DMEM/F12 and RKO in DMEM, all supplemented with 10% fetal bovine serum, 2mM L-Glutamine and 100 U.ml^-1^ penicillin/10μg.ml^-1^ streptomycin. ADO and ATE1 knock-out cells were generated by CRISPR-Cas9 mediated gene editing as described previously^7^. All cell lines were maintained at 37°C in an incubator containing 5% CO_2_. Cells were treated with tert-butyl hydroperoxide (Sigma-Aldrich 416665, 5-6M stock in decane), 2,2-dipyridyl (D216305, 100mM stock in DMSO), MG132 (TargetMol T2154, 25mM stock in DMSO), menadione (Sigma-Aldrich M5625, 10mM stock in DMSO), ICL670A (Cayman Chemicals 16753, 200mM stock in DMSO), ionomycin (Alfa Aesar J62448.M, 1mM stock in DMSO), desferoxamine (DFO, Sigma-Aldrich D9533, 100mM stock in ddH_2_O), bafilomycin A1 (TargetMol T6740, 200μM stock in DMSO), cycloheximide (Merck Life Sciences C4859-1ML, 10mM stock in DMSO), or liproxstatin-1 (TargetMol, 1mM stock in DMSO) through dilution in the culture medium. L/D-alanine powder (Alfa Aesar) was dissolved directly into DMEM/F12 at the desired concentration, then sterile filtered before adding to cells.

### Plasmids and transfections

A pAAV-MCS-mCherry-mDAAO-NES plasmid was kindly provided by Emrah Eroglu (Istanbul Medipol University), and pLentiCMV_HyPer7RgDAAO was a gift from Seth Parker (Addgene plasmid # 217653 ; http://n2t.net/addgene:217653; RRID:Addgene_217653). 1μg of plasmid diluted in 360μL Opti-MEM and 40μL PEI (1mg.ml^-1^) was transfected into SH-SY5Y cells seeded in a 6 cm dish at 70% confluency. 24 hours later, cells were treated with 1μg.mL^-1^ puromycin for 7 days to select for a stably expressing population. SH-SY5Y cells stably expressing doxycycline-inducible, C-terminally V5-tagged RGS4 (RGS4:V5), or a mutant in which the cysteine at position 2 was replaced with an alanine, were generated through transduction using lentiviral particle-containing supernatant produced by transfecting HEK 293T cells with pCW57-RGS4:V5 or pCW57-RGS4(C2A):V5 alongside the viral packaging plasmids pCMVR8.74 and pMD2.G. Transduced cells were selected for by exposure to 1μg.mL^-1^ puromycin for 7 days. RGS4:V5 protein abundance was subsequently induced by treating cells with 1μg.mL^-1^ doxycycline for 24 hours. SH-SY5Y cells stably expressing RGS4_1-11_GFP were generated as described previously^9,21^.

### Immunoblotting

Protein samples were collected in lysis buffer (10 mM Tris pH 7.5, 0.25 M NaCl, 0.5% Igepal) supplemented with Complete^TM^ protease inhibitor cocktail (Sigma Aldrich). Lysates were centrifuged at 13,000rpm for 3 minutes at 4°C, and the supernatant mixed with Laemmli sample buffer. Proteins were then separated via SDS-PAGE electrophoresis. Membranes were blocked in 4% milk for 1 hour, then incubated in primary antibody overnight: HIF-1α (610959, BD Biosciences), RGS4 (15129, CST), RGS5 (sc-514184, SCBT) IL-32 (sc-517408, SCBT), ADO (ab134102, Abcam), ATE1 (HPA038444,

Human Protein Atlas) LC3 (L8918, Sigma), V5 (SV5-P-K, Chromotek), and GFP (11814460001, Sigma Aldrich). HRP-conjugated secondary antibodies were sourced from DAKO and used in conjunction with chemiluminescence substrate (West Dura, 34076, Thermo Fisher Scientific) to visualise protein expression using a ChemiDoc XRS+ imaging system (BioRad). β-actin primary antibody was conjugated directly to HRP (ab49900, Abcam). Densitometric analysis was performed using ImageJ software (NIH) and values presented relative to β-actin.

### RT-qPCR

Tri-Reagent (T9424, Sigma Aldrich) was used to extract RNA by phase separation, and cDNA synthesis was performed using equal quantities of RNA using the High-Capacity cDNA Kit (Applied Biosystems). qPCR analysis was performed using Fast SYBR Green Master Mix on a StepOne thermocycler (Thermo Fisher Scientific) using the ΔCt method. The housekeeping gene Hypoxanthine-guanine phosphoribosyl transferase (HPRT) was used as a reference. Sequences for the primers used are as follows;

RGS4 (F_GCAAAGGGCTTGCAGGTCT, R_CAGCAGGAAACCTAGCCGAT),

RGS5 (F_TGGTGACCTTGTCATTCCG, R_TTGTTCTGCAGGAGTTTGT),

xCT (F_ GGGAAAGTCTTGGAACTCAGG, R_ CCAAGTTAGGGATTTAGCTGGTC),

HPRT (F_GACCAGTCAACAGGGGACAT, R_AACACTTCGTGGGGTCCTTTTC).

### Cell viability assay

Cells were seeded at 10,000 cells/well in clear 96-well plates and allowed to grow to confluence, before being treated with increasing concentrations of compound for 4 hours, after which time cells were incubated with MTT (0.5 mg.ml^-1^) for a further 2 hours and absorbance monitored at 562nm using a FLUOstar plate reader (BMG Labtech).

### Measurement of intracellular H_2_O_2_ production from mDAAO

SH-SY5Y cells stably expressing DAAO-P2A-HyPer7 were seeded onto 24mm coverslips and allowed to reach 70% confluency. Coverslips were then mounted in an imaging chamber (AttoFluor, Thermo Fisher) and placed on the temperature-controlled stage (set to 37°C) of an inverted, widefield fluorescence microscope (Leica DMi8). Cells were maintained in an imaging buffer containing; 25 mM HEPES, 117 mM NaCl, 1 mM NaH_2_PO_4_, 11 mM D-glucose, 4.5 mM KCl, 1 mM MgCl_2_ and 1mM CaCl_2_. L/D-alanine or tBHP (as 25X solutions) were added dropwise to the recording bath as indicated. Cells were illuminated using an LED light source (pE-340, CoolLED) passed sequentially through either a cyan (excitation 405 ± 20 nm, 450nm dichroic mirror, emission filter 500 ± 20 nm) or GFP (excitation filter 470 ± 20 nm, 495 nm dichroic mirror, emission filter 525 ± 25 nm) filter cube. Images were captured every 3 seconds using a 63x objective (HC PL FLUOTAR 63x/1.30 OIL, Leica Microsystems CMS) and a monochrome camera (Prime 95B, Photometrics). Raw images were background subtracted using a cell-free ROI, then the mean fluorescence intensity per ROI in the green channel was divided by the equivalent cyan image to obtain the HyPer7 ratio.

### Lysosomal pH assay

Lysosomal pH was monitored using the Lysosomal Acidic pH Detection Kit (L266 DONJIDO). SH-SY5Y cells were seeded at 40,000 cells per well of an 8-well imaging plate (Ibidi) and allowed to reach 80% confluency. Cells were then washed twice with PBS and incubated in serum-free DMEM/F12 medium containing LysoPrime Green (1:2000) for 30 minutes at 37°C. Cells were then washed again and treated with tBHP/Bafilomycin A1 and pHLys Red (1:1000) for 4 hours. Cells were subsequently washed, stained with Hoechst 33342 (10μg.mL^-1^, H1399 Thermo fisher) and imaged using a DMi8 widefield microscope using DAPI (excitation 350 ± 25 nm, 400 nm dichroic mirror, emission 460 ± 25 nm), GFP (excitation 470 ± 20 nm, 495 nm dichroic mirror, emission 525 ± 25 nm) and rhodamine (excitation 546 ± 5 nm, 560 nm dichroic mirror, emission 585 ± 20 nm) filter cubes. Images were taken using a 20x air immersion objective (HC PL FLUOTAR 20x/0.55). Images were quantified by using the green channel to generate a signal-based mask, with the inverted mask used for background subtraction. A ratio was calculated by dividing the mean red fluorescence intensity by the green fluorescence intensity within the masked regions.

### Lipid peroxidation assay

To monitor lipid peroxidation, SH-SY5Y cells were seeded at 40,000 cells per well of an 8-well imaging plate (Ibidi) and allowed to reach 80% confluency. Cells were then incubated in serum-free DMEM/F12 medium containing 2.5μg.mL^-1^ BODIPY-C11 (D3861, Thermo Fisher) for 30 minutes at 37°C. The unincorporated BODIPY-C11 was washed off with PBS then cells were treated with tBHP/Liproxstatin-1 for 4 hours in Fluorobrite medium (A1896701, Thermo Fisher) with 1% FBS. Cells were subsequently stained with Hoechst 33342 (10μg.mL^-1^) and imaged as described above, using a 63x objective. For each image, the red channel was used to generate a signal-based mask and the inverted mask was used for background subtraction. A ratio was then calculated by dividing the mean green fluorescence intensity by the corresponding red fluorescence intensity within the masked regions.

## Acknowledgments and funding sources

This work was funded by the Wellcome Trust (301530/Z/23/Z) and the British Heart Foundation (FS/IBSRF/24/25194), with support from the BHF Oxford Centre of Research Excellence, University of Oxford and the Oxford Branch of the Ludwig Institute for Cancer Research (PJR, Distinguished Scholar). This work was also supported by the Francis Crick Institute, which receives its core funding from Cancer Research UK (FC001501), the UK Medical Research Council (FC001501) and the Wellcome Trust (FC001501).

## Notes

### Competing Interest Statement

The authors have declared no competing interest.

### Summary of Updates

Revisions to figures 1 and 4, author list updated

